# The Impact of Storage Condition and Duration on Function of Native and Cargo-Loaded Mesenchymal Stromal Cell Extracellular Vesicles

**DOI:** 10.1101/2022.06.14.496108

**Authors:** Daniel Levy, Anjana Jeyaram, Louis J. Born, Kai-Hua Chang, Sanaz Nourmohammadi Abadchi, Angela Ting Wei Hsu, Talia Solomon, Amaya Aranda, Samantha Stewart, Xiaoming He, John W. Harmon, Steven M. Jay

**Affiliations:** Fischell Department of Bioengineering, University of Maryland, College Park, MD 20742, USA; Department of Surgery, Johns Hopkins University School of Medicine, Baltimore, MD 21224, USA; Program in Molecular and Cell Biology, University of Maryland, College Park, MD 20742, USA

**Author notes:** **Correspondence**: Steven M. Jay, Fischell Department of Bioengineering, University of Maryland, 3116 A. James Clark Hall, College Park, MD 20742. These authors contributed equally to this work (Daniel Levy, Anjana Jeyaram).

**Keywords:** Exosomes, microvesicles, miR-146a-5p, HOTAIR

## Abstract

As evidenced by ongoing clinical trials and increased activity in the commercial sector, extracellular vesicle (EV)-based therapies have begun the transition from bench to bedside. As this progression continues, one critical aspect to EV clinical translation is understanding the effects of storage and transport conditions. Several studies have assessed the impact of storage on EV characteristics such as morphology, uptake, and component content, but effects of storage duration and temperature on EV functional bioactivity and, especially, loaded cargo are scarcely reported. Here, EV outcomes following storage at different temperatures (room temperature, 4°C, -20°C, -80°C) for various durations as well as after lyophilization were assessed. Mesenchymal stem/stromal cell (MSC) EVs were observed to retain key aspects of their bioactivity (pro-vascularization, anti-inflammation) for up to 4-6 weeks at -20°C, -80°C, and after lyophilization. Furthermore, via *in vitro* assays and an *in vivo* wound healing model respectively, these same storage conditions were also demonstrated to enable preservation of the functionality of loaded microRNA (miRNA) as well as long non-coding RNA (lncRNA) cargo in MSC EVs. These findings extend the current understanding of how EV therapeutic potential is impacted by storage conditions and may inform best practices for handling and storage of MSC EVs for both basic research and translational purposes.

## 1. Background

Extracellular vesicles (EVs) are nanoscale cell-derived products that are implicated as promising therapeutic agents and drug delivery vehicles [1]. From a drug carrier perspective, EVs compare favorably to synthetic delivery systems with respect to cargo delivery efficiency, physiological transport properties, and multifunctionality [2]–[5]. As native therapeutic agents, mesenchymal stem/stromal cell (MSC) EVs especially have been widely reported to be therapeutically useful in a variety of pre-clinical studies based on their anti-inflammatory and pro-angiogenic effects, leading to their use in clinical trials [6]–[8]. Yet, there remain several obstacles to overcome prior to translation and widespread clinical application of MSC EV therapies. Among these is a relative dearth of knowledge about the effects of storage conditions on MSC EV functionality.

As EVs lack many of the complex cellular machinery and organelles that exist in their parental cells, they may potentially be stored at more desirable conditions for clinical translation such as at -20°C or -80°C freezer storage at room temperature following lyophilization [9]. There have been some important initial studies in this area, specifically into the morphological stability and enzymatic stability of EV cargos as well as towards the determination of appropriate buffers for EV storage [10]–[22]. Further, some knowledge about the effects of storage conditions on EV bioactivity has begun to accumulate. Wu et al. recently demonstrated that cold storage of isolated bEnd.3 EVs at -20°C and -80°C over 28 days resulted in improved cellular uptake and *in vivo* circulation [23], while van de Wakker and colleagues showed that cardiac progenitor cell (CPC)-derived EVs retain their bioactivity in an endothelial cell gap closure assay after 7 days of storage at -80°C, with activity reduced over the same storage period at 4°C [24]. However, despite these and other reports, much remains to be learned about how MSC EV function is impacted by storage.

Here, we add to the understanding of storage effects on EV activity with a focus on human bone marrow-derived MSC EVs. Specifically, our data show that the anti-inflammatory and pro-angiogenic effects of MSC EVs are retained for up to 28 days after storage at -20°C, -80°C and at RT following lyophilization. Furthermore, given the interest in EVs as drug delivery vehicles [25], we examined the effects of storage conditions on loaded microRNA (miRNA) cargo in MSC EVs, showing that while the total amount of cargo detected is decreased in all conditions tested, cargo-associated bioactivity was retained in several storage modes. Lastly, we demonstrate that lyophilization is suitable for preservation of regenerative bioactivity of MSC EVs loaded with the long non-coding RNA (lncRNA) HOTAIR in a db/db mouse wound healing model. Overall, these studies provide further guidance for the field on how EVs, and in particular drug-loaded MSC EVs, should be stored prior to therapeutic application.

## 2. Methods

### 2.1 Cell culture

Human bone marrow-derived MSCs (BDMSCs; ATCC PC-500-012) were cultured in Dulbecco’s Modified Eagle Medium (DMEM) [+] 4.5 g/L glucose, L-glutamine, sodium pyruvate supplemented with 10% EV-depleted FBS, 1% penicillin-streptomycin, and 1% Non-essential amino acids (NEAA) in T175 polystyrene tissue culture flasks. 10% EV-depleted FBS was generated by centrifugation of FBS at 100,000 x g for 16 hours before collection of the non-pelleted supernatant. BDMSCs were grown until passage 3 or 4 before EV isolation for functional assays and passage 5 for EV characterization. Human umbilical cord vein endothelial cells (HUVECs; Lonza, C2519A) were cultured in tissue culture flasks coated with 0.1% gelatin prior to cell seeding in endothelial growth media (PromoCell, C-C22121). RAW264.7 (ATCC, TIB71) cells were cultured in DMEM [+] 4.5 g/L glucose, L-glutamine, sodium pyruvate supplemented with 1% penicillin-streptomycin and 5% FBS.

### 2.2 EV separation

BDMSCs were grown in EV-depleted media for 2 days before the conditioned medium was collected and subjected to differential ultracentrifugation with a 100,000 x g final centrifugation step as previously described [26]. Pelleted EVs were resuspended in 1x PBS and subsequently washed using Nanosep 300 kDa MWCO spin columns (Pall Laboratories; OD300C35). The washed EVs were then resuspended in 1x PBS and sterile filtered using a 0.2 µm syringe filter.

### 2.3 EV storage conditions

Post-isolation, EVs were aliquoted and stored at either room temperature (RT), 4°C, -20°C, or -80°C into polypropylene tubes. Separate aliquots were prepared for each storage time length (1 week, 4 weeks, 6 weeks) or for freeze-thaw cycles (1 cycle, 5 cycles). Three replicate samples were prepared for each time point and storage condition. Separately, isolated EVs were quantified via total protein amount, aliquoted and lyophilized using a SP VirTis AdVantage Pro Lyophilizer with Intellitronics Controller and stored at RT for the designated storage time (1 week, 4 weeks, 6 weeks). Again, equal amounts of three replicate samples (normalized by protein content) were prepared for each time point.

### 2.4 EV characterization

EVs were quantified by nanoparticle tracking analysis (NTA) using a Nanosight LM10 (Malvern) and NTA analytical software version 2.3. Isolated EVs were diluted to a concentration of ∼10 µg protein/mL and aliquoted for each of the indicated time and storage conditions before loading into the Nanosight analysis chamber at room temperature. Three samples were prepared for each time point. Each sample was analyzed in triplicate using three different fields of view with a 60 second video acquisition time. The camera level and threshold were set at 16 and 7, respectively for all samples.

Total EV protein amount was determined via bicinchoninic acid assay (BCA) following the manufacturer’s protocol. Protein levels were determined with three replicate samples for each time point and each sample was measured in duplicate. Based on the Day 0 protein amount, samples containing 20 µg of protein were aliquoted for each time point for the total protein quantitation and western blots. At each time point, samples were prepared and stored until the blot as run. Total protein stains were done using Swift Membrane Stain (G-biosciences). The membrane was imaged using a LI-COR Odyssey CLX Imager. Specific EV protein marker levels were quantified using western blot analysis for ALIX (Abcam ab186429) at 1:1,000, TSG101 (Abcam ab125011) at 1:1000, CD63 (Applied Biomaterials Inc Y402575) at 1:200, and GAPDH (D16H11; Cell Signaling, 5174) at 1:2000, incubated over two nights at 4°C while shaking. Goat anti-rabbit IRDye 800CW (LICOR 925-32210) secondary antibody was used at a 1:10,000 dilution. The bands were detected with a LI-COR Odyssey CLX Imager and quantified using LI-COR Image Studio.

To obtain TEM images, a negative staining technique was utilized. Isolated EVs were briefly incubated with an aqueous solution of EM grade paraformaldehyde (Electron Microscopy Sciences 157-4-100) before a carbon film grid (Electron Microscopy Sciences CF200-Cu-25) was floated on a droplet of the mixture to incubate. The carbon grid was then washed before floated on a droplet of 1% glutaraldehyde. Again, the grid was washed before floatation on a droplet of uranyl-acetate replacement stain (Electron Microscopy Sciences 22405). Prepared EM grids were then stored before imaging on a JEM 2100 LaB6 TEM.

### 2.5 In vitro bioactivity assays

To assess endothelial gap closure, HUVECs (passage 5) were seeded in 48 well plates in endothelial growth medium (EGM) and allowed to grow until the formation of a confluent monolayer. The cell monolayer was then disrupted to form a scratch using a pipette tip. After washing, the cells were serum-starved for 2 hours after which the medium was replaced with the treatments. EGM and endothelial basal media (EBM) were used as positive and negative controls, respectively. The experiments with Day 0 BDMSC EVs were conducted on the day of isolation and the remaining EVs were stored at the indicated conditions or freeze-thaw cycles. After storage, another gap closure assay was performed with the samples at those time points. The gap area was imaged at both 0 and 11 hours. The change in denuded area was quantified using ImageJ.

Endothelial tube formation was assessed using HUVECs (passage 5). Cells were washed, trypsinized and diluted in EBM supplemented with 0.1% FBS before cell counting. HUVECs were then aliquoted, pelleted at 300 x g, where the supernatant was removed, and the cells were resuspended in their respective treatments of stored BDMSC EVs (100 µg/mL) in EBM. Resuspended HUVECs were then added to Growth Factor Reduced Matrigel (Corning, 356252) coated 24-well plates at a seeding density of 75,000 cells/well. Phase-contrast images of tube-forming HUVECs were then taken after 2-12 hours and the number of branch points was quantified using ImageJ.

To evaluate effects of EVs on IL-6 secretion, RAW264.7 mouse macrophages were seeded into 48 well plates in DMEM supplemented with 5% FBS and 1% Penicillin-streptomycin at a density of 100,000 cells/well. 24 hours post-seeding, RAW264.7s were pre-treated with either no treatment, dexamethasone (Dex; Sigma-Aldrich D4902-25MG), or the given BDMSC EV treatments (100 µg/mL) in cell culture media. 24 hours later, RAW264.7 treatments were removed and replaced with 10 ng/mL LPS (Sigma-Aldrich L4391-1MG) in DMEM supplemented with 5% FBS and 1% Pen-strep for an additional 4 hours. After LPS treatment, cell supernatants were collected and stored at -80°C before assessment via ELISA. IL-6 levels from RAW264.7 conditioned medium was quantified using a DuoSet ELISA kit (R&D systems, DY406) following the manufacturer’s instructions.

### 2.6 EV loading via sonication

The same day as BDMSC EV isolation, EVs were loaded with a miR-146a-5p mimic (Dharmacon, C-300630-03-0050) as previously described [27]. 100 µg of EVs were incubated with 1000 pmol of miR-146a-5p in 100 µL PBS for 30 minutes. EVs were then sonicated in a water bath sonicator (VWR® symphony™; cat# 97043-964, 2.8 L capacity, dimensions 24 L × 14 W × 10 D cm) twice for 30 seconds at 35 Hz, resting on ice for 1 minute between sonications. After sonication, EVs were washed in a 300 kDa MWCO filter with 1x PBS three times. Loaded EVs were then pooled, quantified for total protein content via BCA and aliquoted for storage in their respective conditions.

At the indicated time points and storage conditions, loaded BDMSC EV samples were placed in 700 µL Qiazol lysis reagent (Qiagen, 79306) spiked with 2 fmol cel-miR-39 (Norgen Biotek, 59000) as an internal control. Total RNA was isolated with a miRneasy mini kit (Qiagen, 217004) using the manufacturer’s instructions. Post-RNA isolation, cDNA was generated from isolated total RNA samples using a miScript II RT kit (Qiagen, 218161). cDNA was stored at -20°C before qPCR, which was performed on a QuantStudio 7 Flex qPCR system (ThermoFisher Scientific, 4485701) using SsoAdvanced Universal SYBR (Bio-Rad, 1725271). Transcripts for miR-146a-5p were quantified with cel-miR-39 as the internal control. The PCR primer sequences are listed as follows:

miR-146a-5p FWD:

CGCAGGAGAACTGAATTCCA

miR-146a-5p REV:

CAGGTCCAGTTTTTTTTTTTTTTT

cel-miR-39 FWD:

GTCACCGGGTGATAAATCAG

cel-miR-39 REV:

GGTCCAGTTTTTTTTTTTTTTTCAAG

Transcript expression was calculated using the comparative Ct method normalized to cel-miR-39 (2-ΔΔCt) and expressed as fold-change of miR-146a-5p transcripts in storage groups versus freshly loaded EVs.

### 2.7 Animal studies

P4 BDMSCs were seeded at 500,000 cells/flask (T175) before transfection as previously described with a pCMV-HOTAIR plasmid [26]. Specifically, pCMV-HOTAIR was transfected into BDMSCs via Lipofectamine 3000 (ThermoFisher Scientific, L30000015) for one hour, before washing and growth in EV-depleted media. Conditioned media was then collected on days 3-6 post-transfection and isolated via ultracentrifugation as described above before subsequent lyophilization. Mice (db/db, 40-50 g; Jackson Laboratory) were anesthetized with isoflurane before shaving of their dorsum. An 8 mm punch biopsy was taken from the shaved region of each mouse and buprenorphine (0.05 mg/kg) was subcutaneously injected on day 0, 1 and 3. On day 3, four subcutaneous injections (1 µg/µL; 50 µL) were given in a cross pattern to each mouse. 8 mice were used for each treatment group. Tracing of wounds and wound closure were determined as previously described [26]. All animal experiments were performed in compliance with the National Institutes of Health guide for care and use of laboratory animals.

### 2.8 Statistics

Two-way analysis of variance – ANOVA with Dunnett’s multiple comparison test (compared to Day 0 measurements) were utilized to determine statistical significance in all *in vitro* experiments. Two-way ANOVA with Holm-Sidak’s multiple comparison test was used to determine statistical significance in the *in vivo* wound healing time-course experiments. Statistical analyses were performed with Prism 9 (Graphpad Software). Significance notation in presented figures: ns = *p* > 0.05, \**p* < 0.05, \*\**p* < 0.01, \*\*\**p* < 0.001, \*\*\*\**p* < 0.0001

## 3. Results

### 3.1 Frozen storage and lyophilization lead to increased EV size with little change in EV-associated proteins

EVs were isolated from the conditioned medium of BDMSCs via ultracentrifugation, aliquoted and stored in each of their respective conditions and time points. EV size distribution and concentration were assessed for each storage condition via nanoparticle tracking analysis (NTA) using a NanoSight LM10. Size distributions of isolated EVs were within expected size ranges with a representative distribution of BDMSC EVs (∼128 nm mode size) presented (Figure 1A). In our storage studies, an increase in mode size (113 nm to 143 nm, * p≤0.05) was observed after 28 days of storage at -20°C (Figure 1B). After 5 freeze-thaw cycles at -80°C, there was a non-statistically significant increase in mode size (113 nm to 145 nm) (Figure 1C). The increase in size is postulated to be due to aggregation of EVs during storage, which is supported by an observed increase in mean size as detected via NTA (Supplementary Figure 1). TEM analysis also confirmed that EV morphology was retained after lyophilization (Figure 1D). Further, we did not observe significant changes to EV protein markers CD63 and TSG101 with storage at room temperature (RT), 4°C, -20°C, and -80°C or after up to 5 freeze-thaw cycles as determined via immunoblotting (Figure 1E).

**Figure 1.**
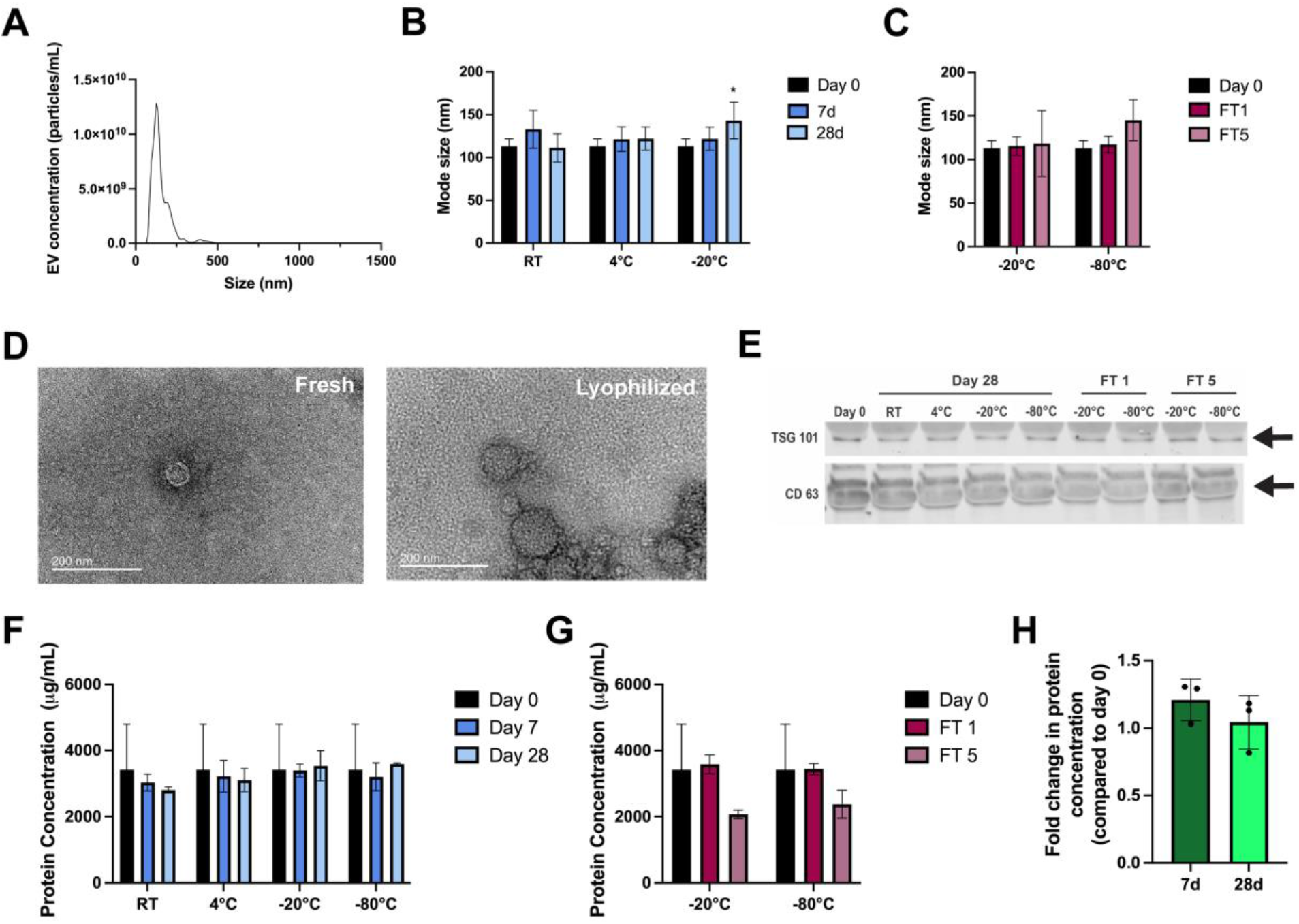
Morphological, size, and protein characterization of stored BDMSC EVs. (A) Representative concentration and size distribution profile of isolated BDMSC EVs as assessed by NTA. (B) Mode size (diameter) changes under varying storage conditions of BDMSC EVs at day 0, 7 and 28. (C) Mode size (diameter) changes of BDMSC EVs after 1 freeze-thaw (FT1) and 5 freeze-thaw cycles (FT5) compared to day 0. (D) TEM images of isolated BDMSC EVs immediately after isolation or after lyophilization and reconstitution. (E) Immunoblot analysis of BDMSC EV markers TSG101 and CD63 after storage at room temperature (RT), 4°C, -20°C, -80C, or freeze-thaw cycles FT1 and FT5. (F) Protein concentrations of BDMSC EVs under varying storage conditions of RT, 4°C, -20°C, or -80°C after 7 or 28 days as compared to day 0. Concentration was determined via BCA assay. (G) Protein concentrations of BDMSC EVs after FT1 or FT5 compared to day 0, assessed via BCA assay. (H) Changes in protein concentration of BDMSC EVs following lyophilization and storage at RT for 7 or 28 days compared to pre-lyophilization. Data are representative of three independent experiments (n=3); All values are expressed as mean ± standard deviation (*p < 0.05, **p < 0.01).

Again, isolated BDMSC EVs were aliquoted and stored in their respective conditions and time points before assessing total protein content using a BCA assay. While storage at -20°C and -80°C generally preserved total protein content, we observed a slight trend of decreasing protein content over up to 28 days when stored at RT and 4°C (Figure 1F). Furthermore, when subjected to freeze-thaw cycles, a non-statistically significant decrease in total protein concentration was observed after 5 freeze-thaw cycles at both -20°C and -80°C (Figure 1G). Total protein content within separate isolated BDMSC EV samples was determined via BCA assay, and aliquots equivalent to 100 µg were lyophilized and stored for either 7 or 28 days before re-assessing total protein content. We observed that lyophilization largely preserves the total protein content of BDMSC EVs (Figure 1H).

### 3.2 Preservation of BDMSC EV in vitro bioactivity is greatly impacted by their storage condition

EVs were isolated from BDMSC conditioned medium via ultracentrifugation, aliquoted and stored for 4 weeks at either RT, 4°C, -20°C, -80°C, or RT after lyophilization. Vascularization bioactivity of the stored BDMSC EVs was then assessed via an endothelial gap closure assay, with freshly isolated BDMSC EVs from the same donor used as a positive control. At a dose of 100 µg/mL, the freshly isolated, -20°C, -80°C, and lyophilized BDMSC EV groups induced an increase in HUVEC gap closure when compared to the negative control, while the RT and 4°C treatment groups resulted in no observable improvement compared to the negative control media condition (Figure 2A). To test whether freeze-thaw cycles have an impact on BDMSC EV bioactivity, EVs were isolated and stored at either -20°C or -80°C and subjected to either 1 or 5 freeze-thaw cycles while in storage for 7 days. In another gap closure assay, it was observed that while 1 freeze-thaw cycle had minimal impacts on gap closure, 5 cycles led to a marginal decrease in gap closure when compared to freshly isolated BDMSC EVs (Figure 2B).

**Figure 2.**
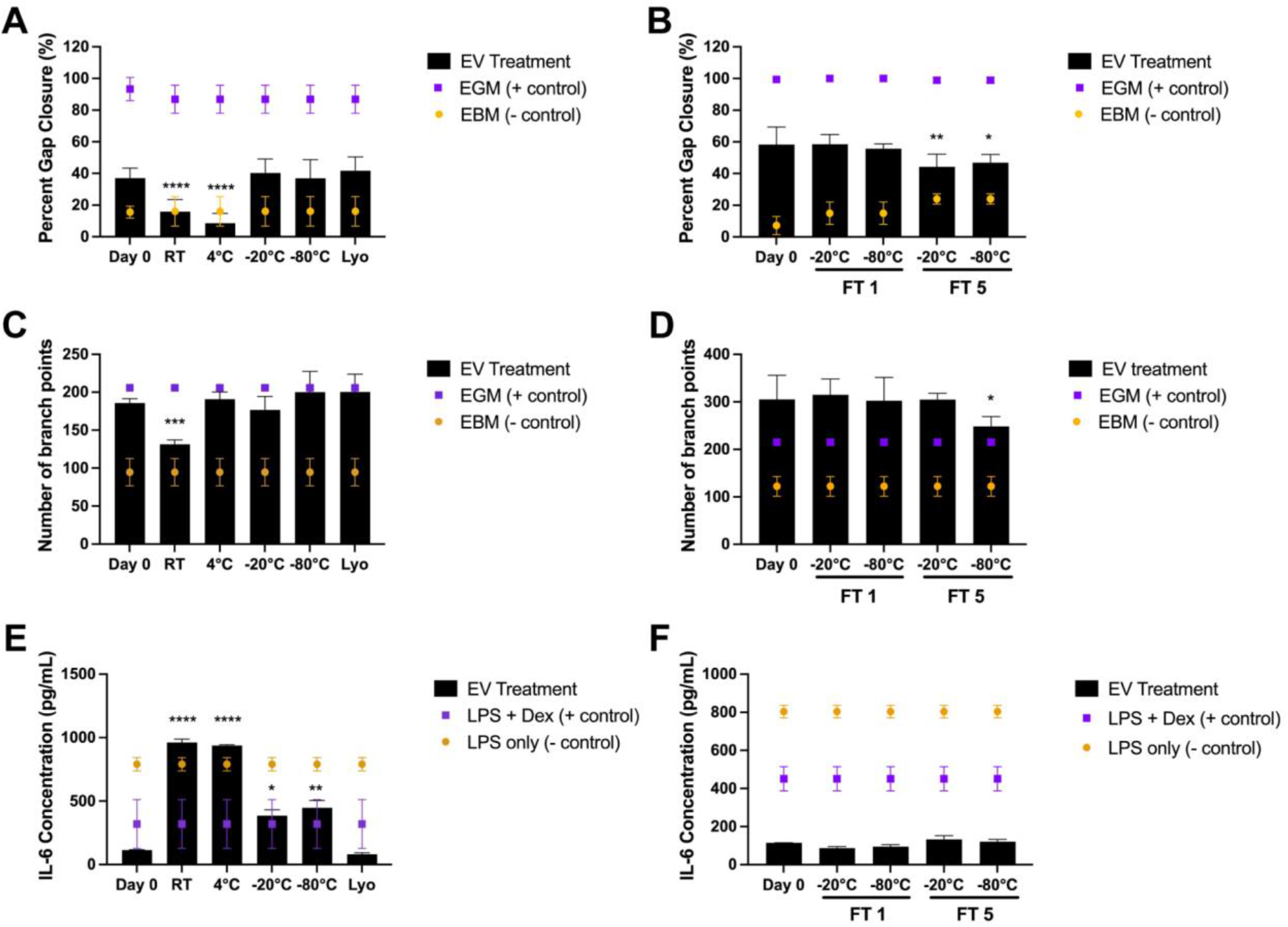
Storage conditions affect in vitro therapeutic EV functionality. (A) Post-scratch induction, HUVECs were treated with either endothelial growth medium (positive control), basal medium (negative control), or 100 µg/mL BDMSC EVs stored for 4 weeks at either RT, 4°C, -20°C, -80°C or lyophilized (Lyo; stored at RT after lyophilization). Changes in gap closure were compared to BDMSC EVs isolated at day 0, before storage. (B) Post-scratch induction, HUVECs were treated with either endothelial growth medium, basal medium, or 100 µg/mL BDMSC EVs after storage at either -20°C or -80°C and FT1 or FT5. Changes of gap closure were compared to BDMSC EVs isolated at day 0, before storage and freeze-thaw. (C) Tube formation capabilities of HUVECs after treatment with BDMSC EVs stored at either RT, 4°C, -20°C, -80°C, or Lyo for 4 weeks. (D) Tube formation capabilities of HUVECs after treatment with BDMSC EVs after storage at either -20°C or -80°C and FT1 or FT5. (E) RAW264.7 macrophages were pre-treated with BDMSC EVs stored at either RT, 4°C, -20°C, -80°C, or Lyo (6 week storage time). RAW264.7s were then treated with LPS before collection of cell supernatant and IL-6 level quantification via ELISA. (F) RAW264.7 macrophages were pre-treated with BDMSC EVs stored at either -20°C or -80°C and FT1 or FT5. Similarly, after LPS stimulation, IL-6 levels in cell supernatants were quantified via ELISA. All values are expressed as mean ± standard deviation (*p < 0.05, **p < 0.01, ***p < 0.001, ****p < 0.0001).

To confirm whether the *in vitro* pro-vascularization potential of BDMSC EVs is impacted by storage conditions, a similar treatment scheme was performed using an endothelial tube formation assay. Again, we observed a significant decrease in bioactivity in the BDMSC EVs stored at RT while -20°C, -80°C and lyophilization preserved bioactivity; interestingly, storage at 4°C also preserved the ability of BDMSC EVs to induce tube formation (Figure 2C). The tube formation assay was also performed using the freeze-thaw groups at -20°C and -80°C and 1 or 5 freeze-thaw cycles. As in the gap closure assay, 1 freeze-thaw cycle had no effect on bioactivity, while a slight decrease was observed after 5 cycles at -80°C (Figure 2D).

To further confirm whether key BDMSC EV bioactivity is impacted by storage conditions, the anti-inflammatory effects of BDMSC EVs were assessed in an LPS-stimulated RAW264.7 mouse macrophage model, with the key output being the amount of pro-inflammatory IL-6 secretion post-LPS stimulation, as this has been shown to correlate with MSC EV anti-inflammatory activity *in vivo* [28]. Using 6 weeks storage time as an endpoint, it was observed that the BDMSC EVs from the RT and 4°C groups lost their ability to attenuate IL-6 secretion compared to freshly isolated BDMSC EVs (Figure 2E). BDMSC EVs stored at -20°C and -80°C lowered IL-6 secretion, but to a lesser extent than freshly isolated EVs. Additional experiments revealed that BDMSC anti-inflammatory activity as measured by this assay was reduced after 4 weeks of storage at 4°C, with continued loss of activity over time, while activity was retained in EVs stored at -20°C and -80°C for up to 8 weeks (Supp. Figure 2). Interestingly, lyophilized BDMSC EVs stored at RT completely preserved the ability to reduce IL-6 levels. It was also confirmed that over the course of 1 week, the impact of freeze-thaw on BDMSC EV bioactivity was negligible in this LPS-stimulated RAW264.7 mouse macrophage model (Figure 2F).

### 3.3 Storage of miRNA-loaded EVs at -20°C, -80°C, and after lyophilization better preserves miRNA content and bioactivity compared to higher temperature storage conditions

A significant part of the therapeutic potential associated with EVs lies in their use as drug carriers. To assess the effects of storage conditions on cargo-loaded EVs, a sonication method previously validated by our group was chosen for loading of miRNA, which are of particular interest with respect to EV-mediated delivery [27]. BDMSC EVs were again isolated using ultracentrifugation before subsequent total protein quantification via BCA assay. Isolated EVs were then promptly co-incubated and sonicated with miR-146a-5p – previously identified as having anti-inflammatory effects – at a ratio of 100 µg to 1000 pmol miRNA as previously described [29]. miR-146a-5p loaded BDMSC EVs were then assessed by NTA (Supplementary Figure 3) and quantified via BCA, aliquoted and stored at either RT, 4°C, -20°C, -80°C, or lyophilized and stored at RT for 4 weeks. After 4 weeks, miR-146a-5p loaded BDMSC EVs samples at each storage condition were again aliquoted and subjected either to RNA isolation, reverse transcription and qPCR or the previously described mouse macrophage inflammatory model. Before RNA isolation and subsequent qPCR, 2 fmol of cel-miR-39 was spiked in as an internal control for the RNA isolation and RT steps. qPCR results demonstrated that storage of loaded EVs led to a marked decrease in miR-146a-5p levels compared to freshly loaded EVs of approximately 99.7%, 95.3%, 80.5%, 81.4%, and 75.1% in loaded EVs stored at RT, 4°C, -20°C, -80°C, or lyophilized and stored at RT respectively (Figure 3A).

**Figure 3.**
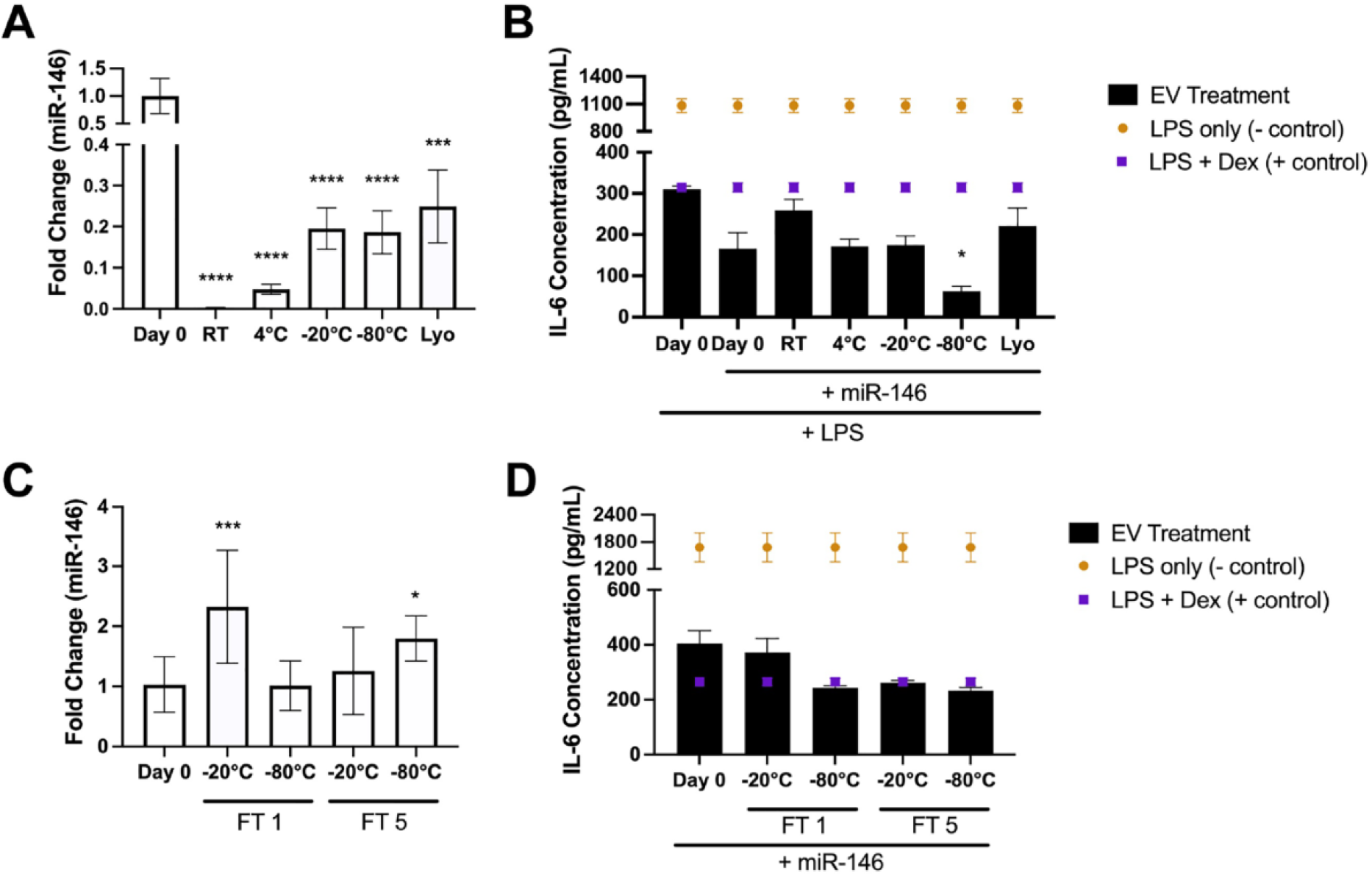
Effects of storage conditions on miRNA content and bioactivity of loaded EVs. (A) Levels of miR-146a-5p in sonicated BDMSC EVs was quantified via qPCR after storage at RT, 4°C, -20°C, -80°C or Lyo after 4 weeks. Fold change of miR-146a-5p was compared to day 0 isolated/loaded BDMSC EVs with 2 fmol cel-miR-39 spiked into samples before RNA isolation as an internal control. (B) RAW264.7 macrophages were again pre-treated with miR-146a-5p loaded BDMSC EVs stored at either 4°C, -20°C, -80°C, or Lyo (4 week storage time). RAW264.7s were treated with LPS before cell supernatants were collected and IL-6 levels were quantified via ELISA. (C) Levels of miR-146a-5p in sonicated BDMSC EVs was quantified via qPCR after storage at either 20°C or -80°C and FT1 or FT5 over 1 week. Again, a spike in of cel-miR-39 was used as an internal control and miR-146a-5p levels were compared to day 0 isolated/loaded BDMSC EVs (D) RAW264.7 macrophages were pre-treated with miR-146a-5p loaded BDMSC EVs stored at either -20°C or -80°C and FT1 or FT5. Similarly, after LPS stimulation, IL-6 levels in cell supernatants were quantified via ELISA. All values are expressed as mean ± standard deviation (*p < 0.05, ***p < 0.001, ****p < 0.0001).

Interestingly, when utilized in the RAW264.7 mouse macrophage inflammatory model at a dose of 40 µg/mL post-sonication, miR-146a-5p loaded BDMSC EVs treatment reduced IL-6 levels at a higher rate than unloaded BDMSC EVs in general; however, changes dependent on storage condition were less pronounced, with only loaded EVs stored at RT leading to significantly reduced anti-inflammatory capabilities (Figure 3B). We also observed that the miR-146a-5p loaded BDMSC EVs stored at -80°C reduced IL-6 levels further than freshly loaded BDMSC EVs, although this outcome may be explained by slight variation within the RAW264.7 assay itself or during the sonication process.

BDMSC EVs loaded with miR-146a-5p were stored at -20°C and -80°C and subsequently subjected to 1 or 5 freeze-thaw cycles. The analysis of these samples yielded highly variable miR-146a-5p levels as assessed by qPCR (Figure 3C). However, in the stimulated RAW264.7 mouse macrophage assay, no statistical differences between miR-146a-5p loaded BDMSC EVs subjected to freeze thaw and freshly loaded EVs were observed (Figure 3D).

### 3.4 Lyophilization preserves the wound healing bioactivity of enhanced BDMSC EVs in vivo

As lyophilization adequately preserved the pro-angiogenic and anti-inflammatory capabilities of BDMSC EVs *in vitro*, we assessed whether these effects translated to a clinically relevant animal model. Previously, we demonstrated that BDMSC EVs loaded with the lncRNA HOTAIR enhanced wound closure after inducing an 8 mm punch biopsy on the dorsum of db/db mice, whereas unaltered BDMSC EVs did not [26]. Thus, we lyophilized HOTAIR-loaded BDMSC EVs and stored them at RT for 4 weeks, before reconstitution and usage in the same db/db mouse model. We observed that lyophilized HOTAIR-BDMSC EVs (90 ± 8.5%) induced a similar improvement in wound healing to fresh HOTAIR-BDMSC EVs (92.8 ± 7% wound closure) compared to the PBS (81 ± 10%) vehicle control (Figure 4).

**Figure 4.**
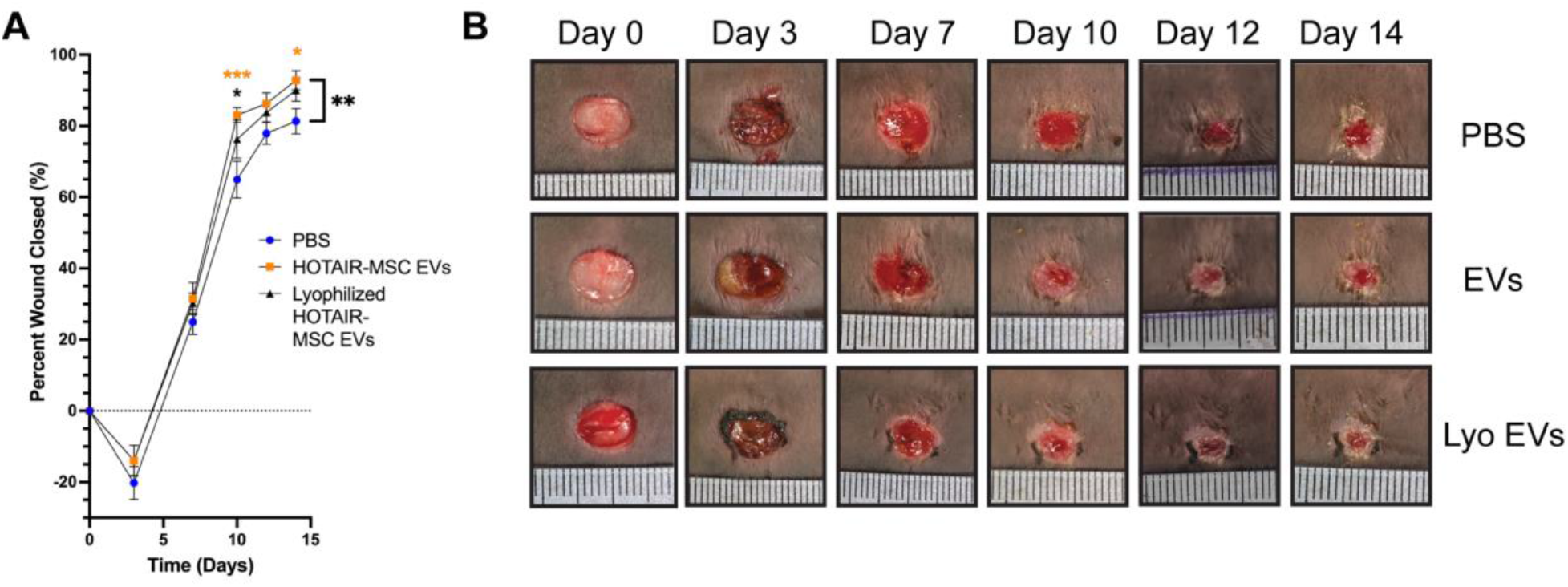
Lyophilization preserves bioactivity of HOTAIR-loaded MSC EVs. Lyophilization and storage at RT for 4 weeks preserved the ability of HOTAIR-loaded BDMSC EVs to accelerate healing in a db/db mouse wound healing model over the course of 2 weeks. (A) Quantification of wound closure as determined by digital planimetry; all values are expressed as mean ± standard deviation (**p < 0.01). (B) Images of wounds at the indicated time points.

## 4. Discussion

EV-based therapies are drawing increasing interest for therapeutic translation for a wide variety of applications [30], [31]. However, to fully realize the clinical potential of EVs, it is vital to increase understanding and optimization of manufacturing parameters such as storage conditions. This reality was reinforced during the rollout of COVID-19 vaccines, with required storage conditions greatly affecting which populations are able to access a given therapeutic [32]. Here, we aimed to determine appropriate storage conditions for MSC EVs, finding that bioactivity of both native and cargo-loaded MSC EVs is affected by storage temperature especially. The data also reinforce the concept that lyophilization can be employed successfully for EV storage. These findings have implications for basic EV research and for eventual clinical translation of EV therapeutics.

Several of our findings are supported by prior work in the field. For example, our results suggest that storage of EVs at -20°C and increased freeze-thaw cycles both lead to EV aggregates, as evidenced by an observed increase in size via NTA (Figure 1B,C). This follows from the report of Wu et al, where a decrease in both total protein content and total RNA content was observed at higher storage temperatures (RT, 4°C), as well as after multiple freeze-thaw cycles [23]. Additionally, our data show that both pro-vascularization and anti-inflammatory bioactivity of BDMSC EVs is diminished after storage at RT and 4°C, while storage at -20°C, -80°C, or lyophilization generally preserves bioactivity to a greater degree for up to 4 weeks (Figure 2). This expands from the work of van de Wakker et al., who showed that CPC EVs exhibit bioactivity in an endothelial cell gap closure assay after 7 days of storage at -80°C, but not 4°C [24].

This work additionally breaks new ground in reporting the effects of storage conditions on cargo-loaded MSC EVs. Using a previously described sonication approach [27] to load BDMSC EVs with the anti-inflammatory miR-146a-5p, we observed that cargo-associated enhanced anti-inflammatory effects were retained in several storage conditions despite an apparent decrease in cargo levels (Figure 3). This could indicate that miR-146a-5p-loaded BDMSC EVs require relatively few copies per EV to achieve profound anti-inflammatory effects, which fits with prior observations that EVs deliver RNA cargo to cells in a highly efficient manner compared to synthetic delivery systems [2]. Also, these results may suggest that the miRNAs that are not tightly bound or associated with EVs are responsible for the majority of the observed miRNA signal in the control group, and that these miRs are removed during the additional washing steps associated with various storage conditions. In this case, additional preparation steps using detergents, enzymes, or other methods to degrade miRs interacting with EV membranes may be useful to determine more accurate cargo loading levels for predicting bioactivity in future studies.

Critically, we observed that lyophilization of HOTAIR-loaded BDMSC EVs, preserved their enhanced bioactivity in a db/db wound healing mouse model (Figure 4), indicating that lyophilization and storage at RT is suitable for retaining the activity of enhanced EVs. This is a vast improvement when compared to cell-based therapies that require liquid nitrogen phase storage during transport and upon receipt in the clinic, further supporting the translatability of EV therapeutics [33].

There are several limitations to the present study that are worth mentioning. Our experiments were limited in duration and do not necessarily account for the full timespan of stability and functionality that would be expected of a clinical EV product. Additional studies over longer time periods are warranted. Further, there are additional considerations beyond what was studied here for fully determining optimal EV preservation conditions. Specifically, studies investigating optimal freezing buffers, vessels, and the use of cryoprotectants to reduce the formation of ice crystals have also been performed previously [11], [15], [34]–[36]. Taken together with the present work, these studies have provided a foundation for determining the optimal storage condition of MSC EVs for preservation of morphology and functionality. Given the current knowledge in the field, and based on our data, we recommend that isolated BDMSC EVs should be stored at either -20°C, -80°C or lyophilized for up to 4 weeks, while minimizing the number of freeze-thaw cycles before either *in vitro* functional bioactivity studies or *in vivo* pre-clinical use.

Overall, our data suggests that both storage condition and duration may have a consequential effect on the therapeutic efficacy of bioactive MSC EVs, including those loaded with therapeutic cargo, in both basic research and clinical translation, with storage at -20°C, -80°C, and lyophilization providing adequate retention of EV activity.

---

**Supplementary Figure 1.**
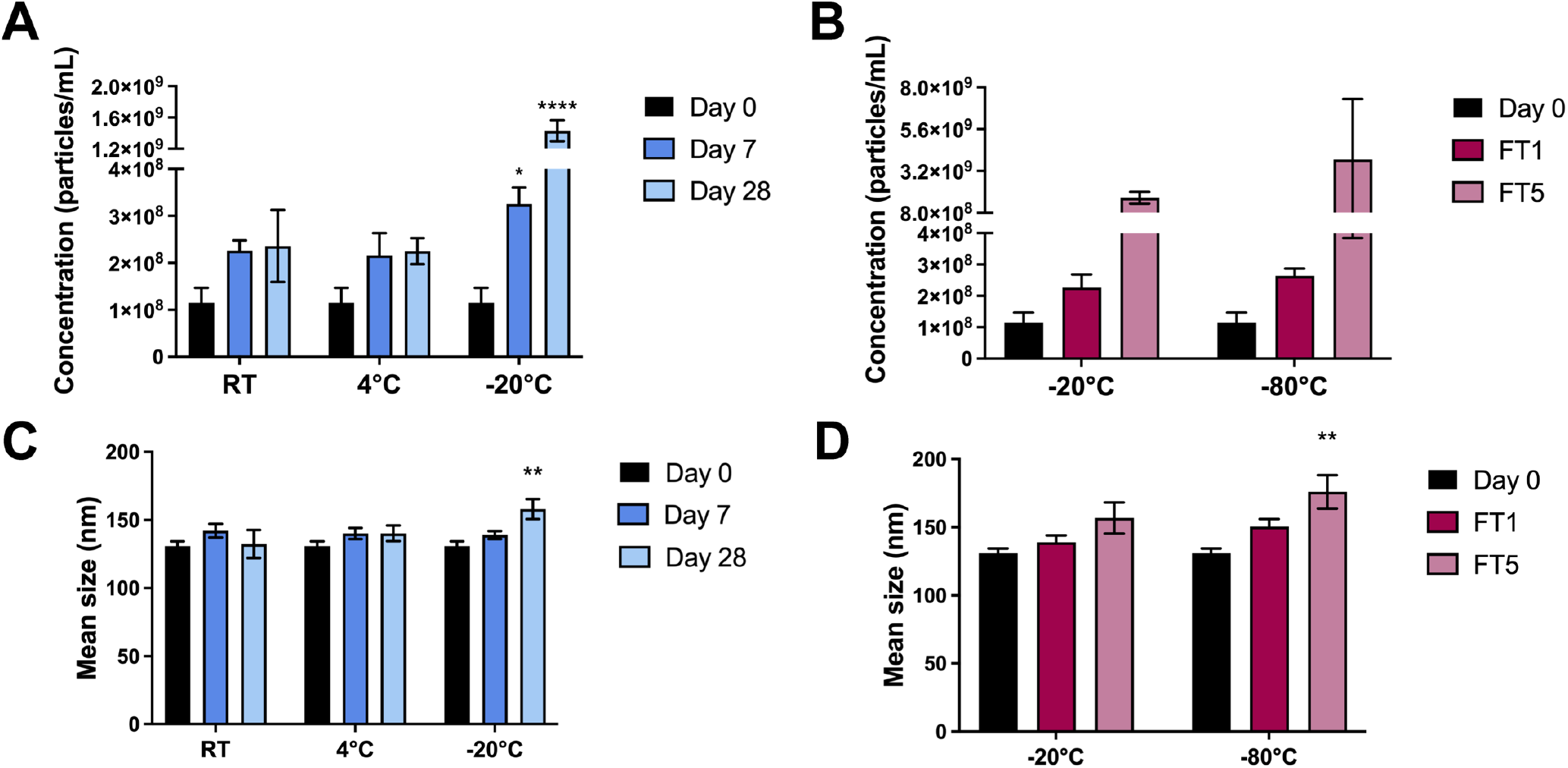
Concentration of BDMSC EV samples after (A) storage up to 28 days and (B) after up to five freeze-thaw cycles. Three separate preparations of BDMSC EVs were isolated and assessed via NTA for particles/mL. (C) Mean size of BDMSC EV samples after storage for up to 28 days and (D) after up to five-freeze thaw cycles. All values are expressed as mean ± standard deviation (*p < 0.05, **p < 0.01, ****p < 0.0001).

**Supplementary Figure 2.**
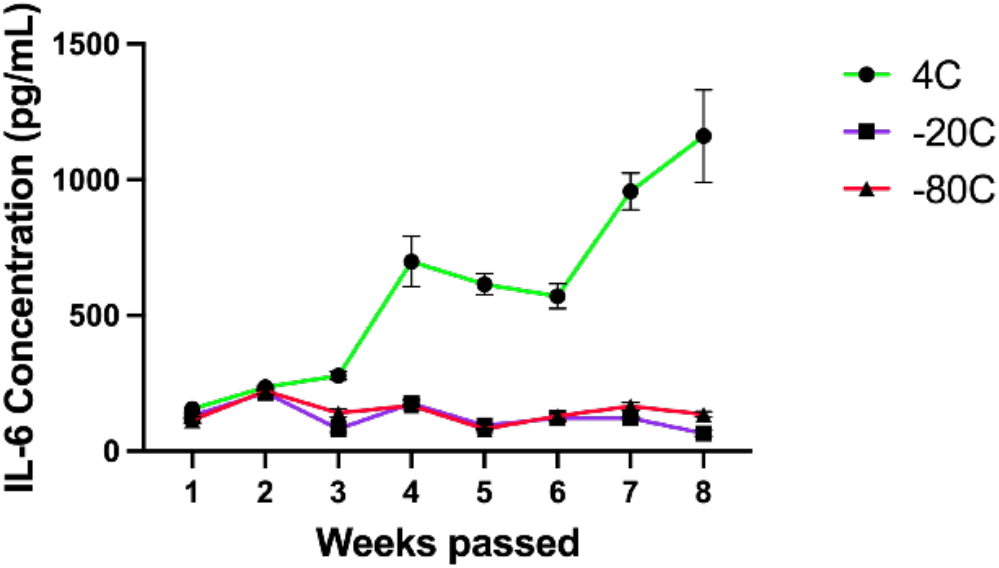
BDMSC EVs (same donor cell source) were isolated weekly for 8 weeks and stored at either 4°C, -20°C, or -80°C before pre-treating RAW264.7. RAW264.7s were then treated with LPS before collection of cell supernatant and IL-6 level quantification via ELISA.

**Supplementary Figure 3.**
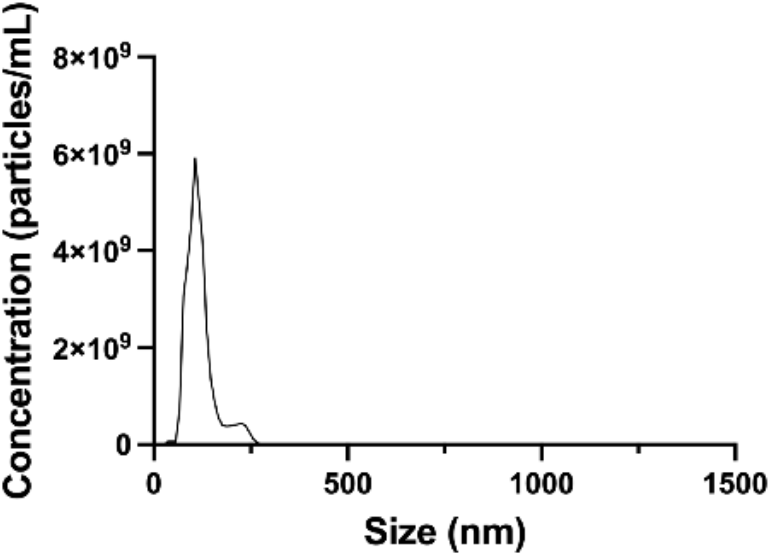
Concentration and size distribution of isolated BDMSC EVs post-sonication with miR-146a-5p as assessed by NTA. Mode size = 113 nm.

