## Supplemental Data for "The Impact of Storage Condition and Duration on Function of Native and Cargo-Loaded Mesenchymal Stromal Cell Extracellular Vesicles"

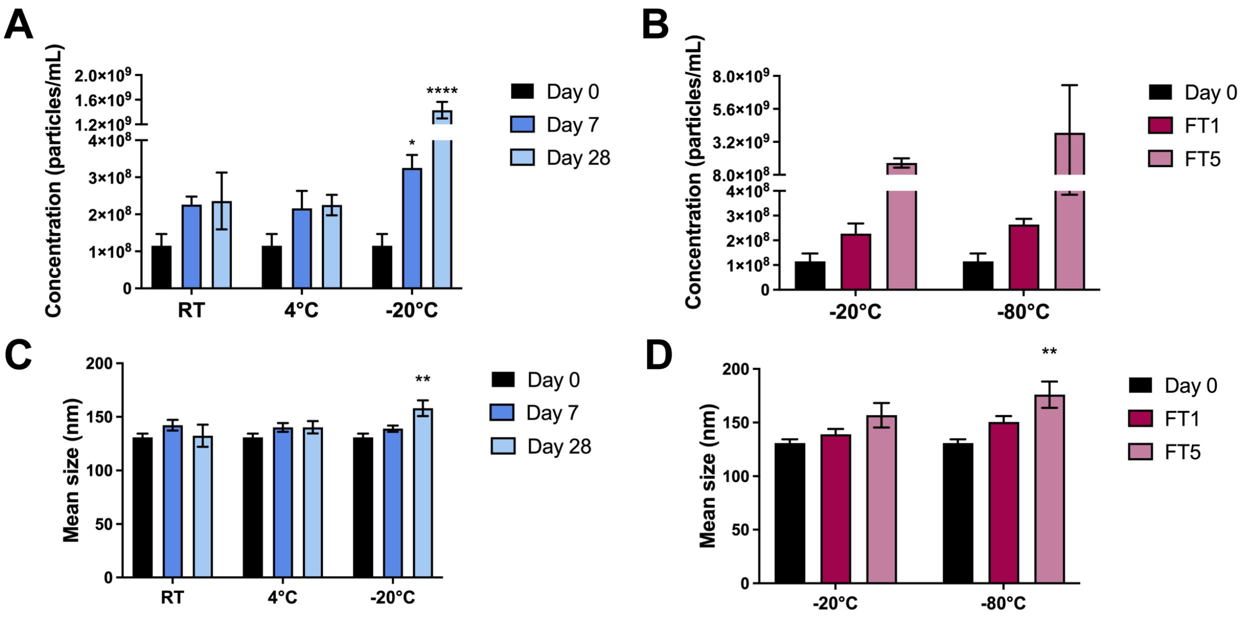


***Supplementary Figure 1.*** Concentration of BDMSC EV samples after (A) storage up to 28 days and (B) after up to five freeze-thaw cycles. Three separate preparations of BDMSC EVs were isolated and assessed via NTA for particles/mL. (C) Mean size of BDMSC EV samples after storage for up to 28 days and (D) after up to five-freeze thaw cycles. All values are expressed as mean ± standard deviation (*p < 0.05, **p < 0.01, ****p < 0.0001).

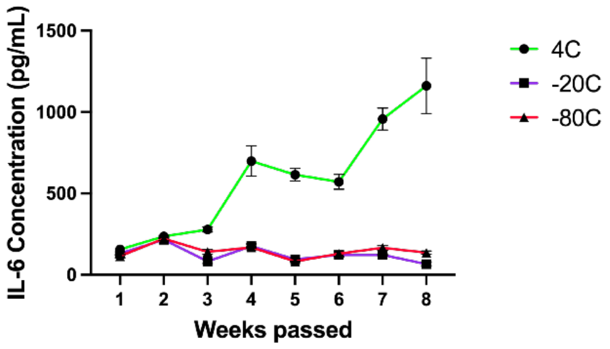


***Supplementary Figure 2.*** BDMSC EVs (same donor cell source) were isolated weekly for 8 weeks and stored at either 4℃, -20℃, or -80℃ before pre-treating RAW264.7. RAW264.7s were then treated with LPS before collection of cell supernatant and IL-6 level quantification via ELISA.

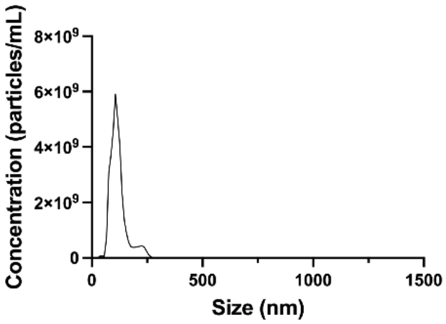


***Supplementary Figure 3.*** Concentration and size distribution of isolated BDMSC EVs post-sonication with miR-146a-5p as assessed by NTA. Mode size = 113 nm.
